# Speech-based identification of L-DOPA ON/OFF state in Parkinson’s Disease subjects

**DOI:** 10.1101/420422

**Authors:** R. Norel, C. Agurto, J.J. Rice, B.K. Ho, G.A. Cecchi

## Abstract

**Background:** Parkinson’s disease patients (PDP) are evaluated using the unified Parkinson’s disease rating scale (UPDRS) to follow the longitudinal course of the disease. UPDRS evaluation is performed by a neurologist, and hence its use is limited in the evaluation of short-term (daily) fluctuations. Subjects taking L-DOPA as part of treatment to reduce symptoms exhibit motor fluctuations as a common complication.

**Objectives:** The aim of the study is to assess the use of speech analysis as a proxy to continuously monitor PDP medication state.

**Methods:** We combine acoustic, prosody, and semantic features to characterize three speech tasks (picture description, reverse counting and diadochokinetic rate) of 25 PDP evaluated under different medication states: “ON” and “OFF” L-DOPA.

**Results:** Classification of medication states using features extracted from audio recordings results in cross-validated accuracy rates of 0.88, 0.84 and 0.71 for the picture description, reverse counting and diadochokinetic rate tasks, respectively. When adding feature selection and semantic features, the accuracy rates increase to 1.00, 0.96 and 0.83 respectively; thus reaching very high classification accuracy on 3 different tasks.

**Conclusions:** We show that speech-based features are highly predictive of medication state. Given that the highest performance was obtained with a very naturalistic task (picture description), our results suggest the feasibility of accurate, non-burdensome and high-frequency monitoring of medication effects.

---

Parkinson’s Disease (PD) is the second most common neurodegenerative disease, after Alzheimer’s. The estimated prevalence of PD in industrialized countries is 0.3% in the general population, 1.0% in people older than 60 years, and 3.0% in people over 80 years old [1]. Around 7 to 10 million people worldwide live with PD; that is more than the combined number of people diagnosed with multiple sclerosis, muscular dystrophy and Lou Gehrig’s disease. In the US, nearly 60,000 Americans are diagnosed with Parkinson’s disease each year, not including the potentially thousands of undetected cases [2]. To follow the progression of PD, the most widely used clinical rating scale is the Unified Parkinson Disease Rating Scale (UPDRS) [3]. It was originally developed in the 1980s and revised in 2001. The UPDRS was not meant for continuous monitoring; it has to be administered by neurologists or motor disorder specialists. As an alternative explored in recent years, speech can potentially be used to monitor patients, to inform on medication effectiveness, and to follow progression. The UPDRS already includes a section for scoring speech in 5 different levels: 0: Normal (no problems); 1: Slight (speech is soft, slurred or uneven); 2: Mild (occasionally parts of the speech are unintelligible); 3: Moderate (frequently parts of the speech are unintelligible); and 4: Severe (speech cannot be understood). However speech scores are inconsistent among graders according to Martinez-Martin *et al.* [4]. Therefore, there is a need for an unbiased measurement of speech changes in the research and clinical communities. Currently the most common treatment of PD includes the use of L-DOPA, which helps ameliorate the symptoms. Unfortunately long-term use of the drug results in symptom control fading out, resulting in fluctuation of medication states known as “ON” and “OFF” states [5, 6].

Speech disorder resulting from neurological impairment such as PD is known as dysarthria. This condition affects mainly the control and execution of movements related to speech production [7]. Previous studies have characterized speech in PD as having the following attributes: reduced loudness, monopitch, monoloudness, reduced stress, breathy and hoarse voice quality, and imprecise articulation [8]. Most recent approaches demonstrate that speech features can help differentiate healthy controls from PDP with high accuracy, in particular using vocal measurements of sustained phonations [9, 10, 11, 12, 13]. There is also evidence of cognitive impairment as part of PD, which affects - or is reflected in - language [14]. Yet another potential difference between PDP and healthy controls or PDP in ON/OFF state is the use of action verbs [15, 16, 17, 18]. Garcia *et al*. [19, 20] and more recently Cotelli *et al*, [21] compared PDP with healthy controls reporting action verb deficit in PDP. Herrera and Cuetos [18] added evidence that PDP without adequate dopamine levels have difficulties in naming action verbs from pictures.

However, there has not been enough research nor strong findings on the evaluation of the use of speech as a way to monitor the medication states [22, 23, 24]. Among those articles, we found that Okada *et al.* [25], analyzed isolated vowel articulation in PD subjects reporting that vowel space area was significantly expanded after L-DOPA treatment contrary to previous findings by [26], where no changes in speech over L-DOPA cycle were found. A recent article [27] analyzes a small cohort of late stage PD subjects using data from a L-DOPA challenge described in [28]. This work, which was limited to the analysis of sustained vowel /a/ and the repetition of a 8-word simple sentence, did not find significant changes in speech.

Therefore, there is a need for objective ways of evaluating PD patients as a way of monitor their disease progression, as well as the effect and duration of their medication. Perhaps there is a prodromal signature in speech that could inform subjects about the risk of PD so they could seek treatment to slow its progression.

In this paper we show that features extracted from speech are informative enough to distinguish the medication state of a PD subject. In particular, we show that a simple and naturalistic task, namely the description of a picture, provides for the highest accuracy, suggesting a potential use in high-frequency and remote monitoring.

## SUBJECTS AND METHODS

### | Subjects

Twenty five subjects (6 females with age of 67 ± 6 years; 19 males with age of 69 ± 7.5 years) with idiopathic Parkinson’s disease. The study was approved by the Tufts University Institutional Review Board.

Inclusion criteria: subjects that respond to L-DOPA treatment, are able to recognize their “wearing off” symptoms, can confirm that they usually improve after their next dose of PD medication, have PD Hoehn & Yahr Stage less than or equal to 3 (assessed while the patient is “ON”), and a score of 26 or more on the Montreal Cognitive Assessment Tool (MoCA) which is the normal range (i.e., no cognitive impairment). Exclusion criteria includes any current history of neurological disease (except for Parkinson’s disease), cognitive impairment, or psychiatric illness that in the investigator’s judgment would interfere with subject participation, treatment with an investigational drug within 30 days, or 5 half lives preceding the first dose of study treatment, whichever is longer, history of regular alcohol consumption exceeding 7 drinks/week for females or 14 drinks/week for males, subjects with cardiac pacemakers, electronic pumps or any other implanted medical devices (including deep brain stimulation devices). Each subject was evaluated by a neurologist using the UPDRS. The differences of speech scores between “OFF” and “ON” states were 2 (1 subject), 1 (7 subjects), 0 (16 subjects), and −1 (1 subject).

### | Design and Protocol

This study reflects the analysis of speech tasks from “Observational Study in Parkinson’s Patient Volunteers to Characterize Digital Signatures Associated with Motor Portion of the UPDRS, daily living activities and speech”, conducted by IBM, Pfizer and Tufts. In this study, three different speech tasks were acquired for each subject at two different sessions: before and after their L-DOPA medication. The order of medication state across sessions was randomized to decrease the learning effect that may influence the results. The first task is called “Picture description” (cookie theft [29, 30] or description of another picture of similar characteristics). In this task, the participant is asked to provide a free-form verbal description of a visual stimulus (the picture). For each session, a different picture was presented to the subject, picture_1 for all subjects in visit_1 and picture_2 for all subjects in visit_2. The second task is a modification of a classic test for mental state evaluation [31, 32] and its called “Reverse counting”. Participants were asked to count in reverse order starting from a different number in each of the sessions to keep the same level of cognitive load. The third task is called “Diadochokinetic rate” and the subjects were asked to pronounce the sequence of three syllables “pa-ta-ka” as rapidly as possible for 10 seconds. This test is widely used for assessing oral motor skills [33].

### | Data Acquisition

The speech tasks were recorded using Audacity software [34] in two channels, one for the analyzed subject and one for the experimenter. The recording parameters were set to sampling frequency of 44.1 kHz with 16 bits and were saved using ‘wav’ format. Both the subject and the experimenter had to wear a headset with a low-impedance unidirectional dynamic microphone.

### | Feature Extraction

The characterization of the speech recordings is performed using two software tools: Python [35] and Praat [36, 37]. Three types of features, which are explained below, were extracted to find signatures of different medication states in the three speech tasks.

### | Mel Frequency Cepstral Coefficients (MFCCs)

Thirteen MFCCs were calculated using python_speech_features package [38]. Following common practice [39], the first coefficient was replaced by the log of the total frame energy in order to analyze the overall energy in the speech of the speaker. The parameters used to calculate the coefficients were windows of 25 ms and windows overlap of 10 ms. To only characterize the voice of the subject, pauses were removed from the recording. A pause is defined by a silence threshold of −25 dB and minimum duration of 100 ms as recommended by Griffiths [40]. To represent the distribution of each coefficient, we computed 10 statistical descriptors, mean, variance, kurtosis, skewness, mode, percentiles 10^th^, 25^th^, 50^th^, 75^th^, and 90^th^. These features are used for all analyzed speech tasks.

### | Nuclei syllable (NS)

As a proxy for fluency in speech (or speech rate), we used the method proposed by De Jong *et al.*[41], to estimate the location of syllables. After locating the syllables, we computed the time lapse between them in the speech recording with and without pauses being removed. To represent the distribution of syllable duration, we computed the following statistical descriptors: mean, variance, kurtosis, skewness, mode, interquartile range (IQR, a measure of variability), 10^th^ percentile, and 90^th^ percentile for a total of 16 features. These features are calculated for the three speech tasks.

### | Semantic features (SF)

Since the picture description task uses free speech, we analyzed the semantic content of the description provided by the subjects. After manually transcribing the recordings, we used the Stanford POS tagger [42] to get part of speech tags for words uttered by subjects. For nouns and verbs we computed the similarity distance to the following seed words: *action, act, move, play, energetic, inaction, sleep, rest, sit* and *wait*. The choice of words was aimed to check whether the use of action vs. non action words was influenced by dopamine level (“ON” vs. “OFF” state of L-DOPA). Words are represented using Gloval vectors for word representations (GloVe) [43]. Briefly, GloVe is an unsupervised learning algorithm for obtaining vector representations for words. Training is performed on aggregated global word-word co-occurrence statistics from a corpus, and the resulting representations showcase interesting linear substructures of the word vector space. We used GloVe version 1.2, using vector representation of 300 dimensions, trained on six billion word corpus taken from Wikipedia 2015 and Gigaword 5 [44]. The distance similarity is computed between each verb and noun uttered by the subject and each of the seed words. To represent the distribution obtained for each seed words, we calculated the following statistical descriptors from the distances of the subjects words: median, 10^th^ percentile, 90^th^ percentile, skewness, kurtosis, IQR. As an additional feature, we also compute the total number of words used in the analysis.

### | Statistical Analysis

Features are ranked based on p-values obtained after performing a two sample t-test. This procedure is applied separately for the training sets generated in the validation procedure. To get an insight of how the features interact in each medication state, we also computed the partial correlations among the top features for each speech task. Partial correlation captures the pattern of covariation between a pair of features by removing the effect of the other analyzed features.

### | Classification

We evaluate the potential of our features to differentiate one medication state from another by applying different classifiers. We chose 4 different classifiers: Nearest Neighbors (NN), Logistic Regression (LR) with l_1_-norm regularization, SVM with elastic regularization and Random Forests (RF). To avoid gender, age and education level confounding effect, we use each subject as his/her own baseline. This means that instead of performing “ON” vs. “OFF” classification, we performed (“ON” - “OFF”) vs (“OFF” - “ON”) classification. Before providing the features to the classifiers, the features are standardized (mean = 0 and standard deviation = 1). Finally, accuracy rates are calculated using a two nested cross-validation approach and we provide their 95% confidence intervals (CI) after performing bootstrap with 1000 samples.

### | Impact of Audio Duration

To evaluate whether a longer recording can provide more informative features, we analyze subjects with recordings of more than 20 seconds in the picture description task (after removing pauses in “ON” and “OFF” states). For each recording, we extract windows of 5, 10, 15, 18 and 20 seconds centered at half of the recording and calculate only MFCC features. To evaluate the impact of the length of the recording on the accuracy rate, we perform 4-fold cross validation based on 100 different data partitions.

## RESULTS

### | Statistical analysis

Table 1 shows the top 5 features for each speech task, where only acoustic features, specifically MFCC #1 (total energy) and MFCC #11, appear to be the most relevant for reverse counting and diadochokinetic rate tasks, respectively. On the other hand top-ranked features for picture description shows SF, NS and MFCC features. For this reason, we only evaluated patterns of co-variation using partial correlation among the top features for the picture description task, which are shown in Fig. 1. It can be seen that the “OFF” state is characterized by a strong positive partial correlations between SF (*play*) and acoustic (MFCC #2) and SF (*act* and *play*) and NS features.

**FIGURE 1.**
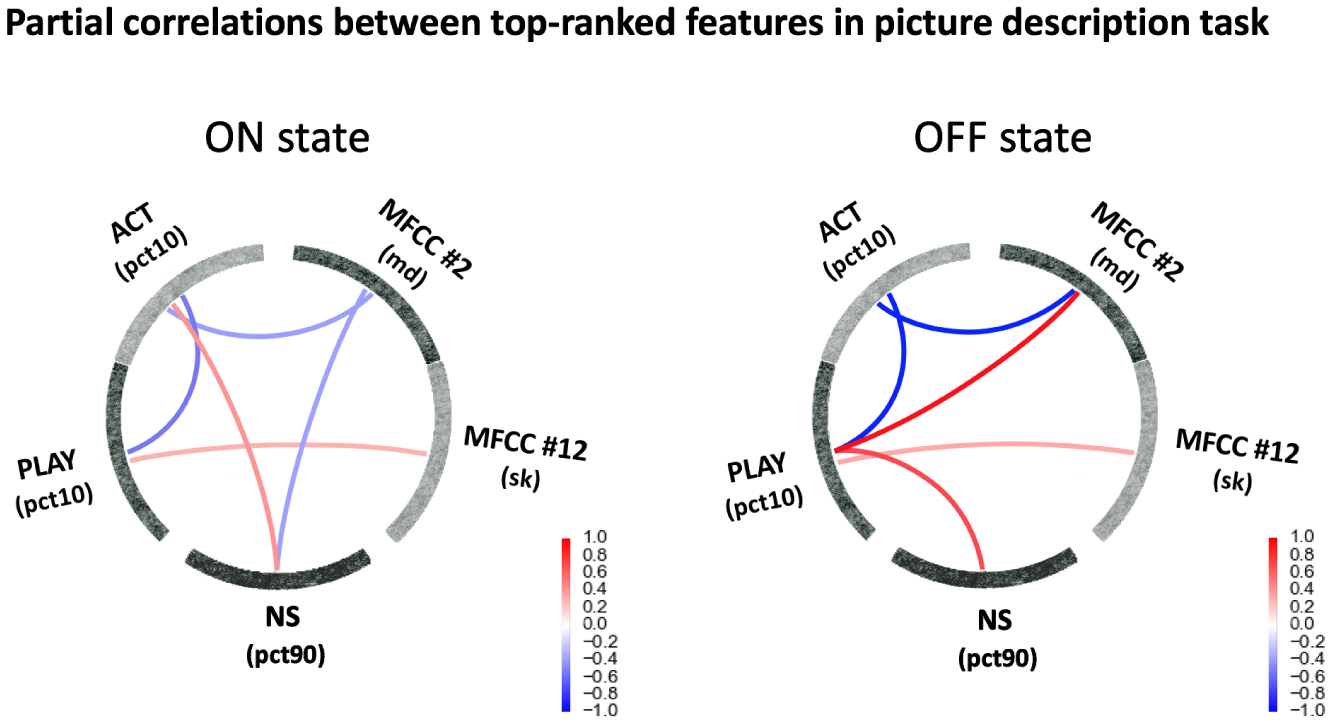
Partial correlations for “ON” and “OFF” states were calculated using the top 5 features of the picture description task described in Table 1. Positive correlations are displayed in red color while negative correlations are in blue. “OFF” state shows a stronger correlation among features in comparison with “ON” state. In addition, new positive correlations were found between SF features (play), NS and MFCC #2

**TABLE 1.** Five top-ranked features for “ON” vs “OFF” states characterization for each speech task. Ranking is calculated with all of the extracted features using two-sample t-test in each training set of our leave-subject-out cross validation approach. Listed statistics are estimated in all samples for reference only. A positive t-statistic indicates greater mean value for the ON state.

| Speech Task | Feature | p-value | t-statistic |
| --- | --- | --- | --- |
| Picture description | PLAY (pct10) | 5.92e-07 | 5.76 |
|  | MFCC #2 (md) | 1.50e-05 | 4.82 |
|  | ACT (pct10) | 2.79e-05 | 4.63 |
|  | MFCC #12 (sk) | 3.96e-04 | -3.81 |
|  | NS (pct90) | 5.10e-04 | 3.73 |
| Reverse counting | MFCC #1 (q50) | 1.54e-07 | -6.14 |
|  | MFCC #1 (q25) | 9.04e-07 | -5.63 |
|  | MFCC #1 (mn) | 4.09e-06 | -5.20 |
|  | MFCC #1 (q75) | 4.88e-06 | -5.15 |
|  | MFCC #8 (sk) | 3.11e-05 | 4.60 |
| Diadochokinetic rate | MFCC #11 (pct75) | 2.46e-05 | 4.69 |
|  | MFCC #11 (mn) | 1.06e-04 | 4.24 |
|  | MFCC #11 (pct50) | 1.86e-04 | 4.06 |
|  | MFCC #3 (pct75) | 2.57e-03 | -3.19 |
|  | MFCC #11 (pct25) | 2.74e-03 | 3.17 |

### | Classification

As explained above, classification tasks are performed by subtracting features from one state to another state. Table 2 shows the classification accuracy using all features and feature selection for the different possible combinations of features in each speech task. LR with l_1_-norm regularization is the classifier that helps to improved performance for several feature combinations. Fig. 2 shows the best performance among the combinations of features (highlighted in Table 2). Results using ten-fold cross-validation are indicated in the barplots. The CI is shown, and all achieve results higher than chance probability.

**TABLE 2.** Performance achieved in each task for different combinations of features. Accuracy rates are calculated with and without feature selection. Only the classifiers with highest accuracy value are shown. The highest accuracy rate is obtained for picture description with and without feature selection. MFCC features are relevant for achieving good performance in the different speech tasks. LR - l_1_-norm achieves improved performance for several combinations possibly due to the sparse nature of the features

| Speech Task<br>(number of subjects) | Features | Accuracy using<br>all features | Classifier for<br>all features | Best accuracy<br>(# features) | Classifier for<br>best accuracy |
| --- | --- | --- | --- | --- | --- |
| Picture description<br>(25 subjects) | NS | 0.64 (16) | NN | 0.76 (15) | NN |
| | SF | 0.68 (61) | LR - $l_1$-norm | 0.84 (1) | NN |
| | NS + SF | 0.80 (77) | LR - $l_1$-norm | 0.84 (1) | NN |
| | MFCC | 0.88 (130) | LR - $l_1$-norm | 0.88 (130) | LR - $l_1$-norm |
| | MFCC + NS | 0.80 (146) | LR - $l_1$-norm | 0.84 (27) | LR - $l_1$-norm |
| | MFCC + SF | 0.76 (191) | LR - $l_1$-norm | 1.0 (5) | LR - $l_1$-norm |
| | MFCC + SF + NS | 0.72 (207) | LR - $l_1$-norm | 1.0 (5) | LR - $l_1$-norm |
| Reverse counting<br>(25 subjects) | NS | 0.76 (16) | NN | 0.76 (16) | NN |
|  | MFCC | 0.76 (130) | RF | 0.88 (82) | NN |
| | MFCC + NS | 0.84 (146) | SVM - elastic | 0.96 (15) | LR - $l_1$-norm |
| Diadochokinetic rate<br>(24 subjects) | NS | 0.71 (16) | RF | 0.71 (15) | RF |
| | MFCC | 0.67 (130) | LR - $l_1$-norm | 0.83 (1) | NN |
|  | MFCC + NS | 0.63 (146) | SVM - elastic | 0.83 (1) | NN |

**FIGURE 2.**
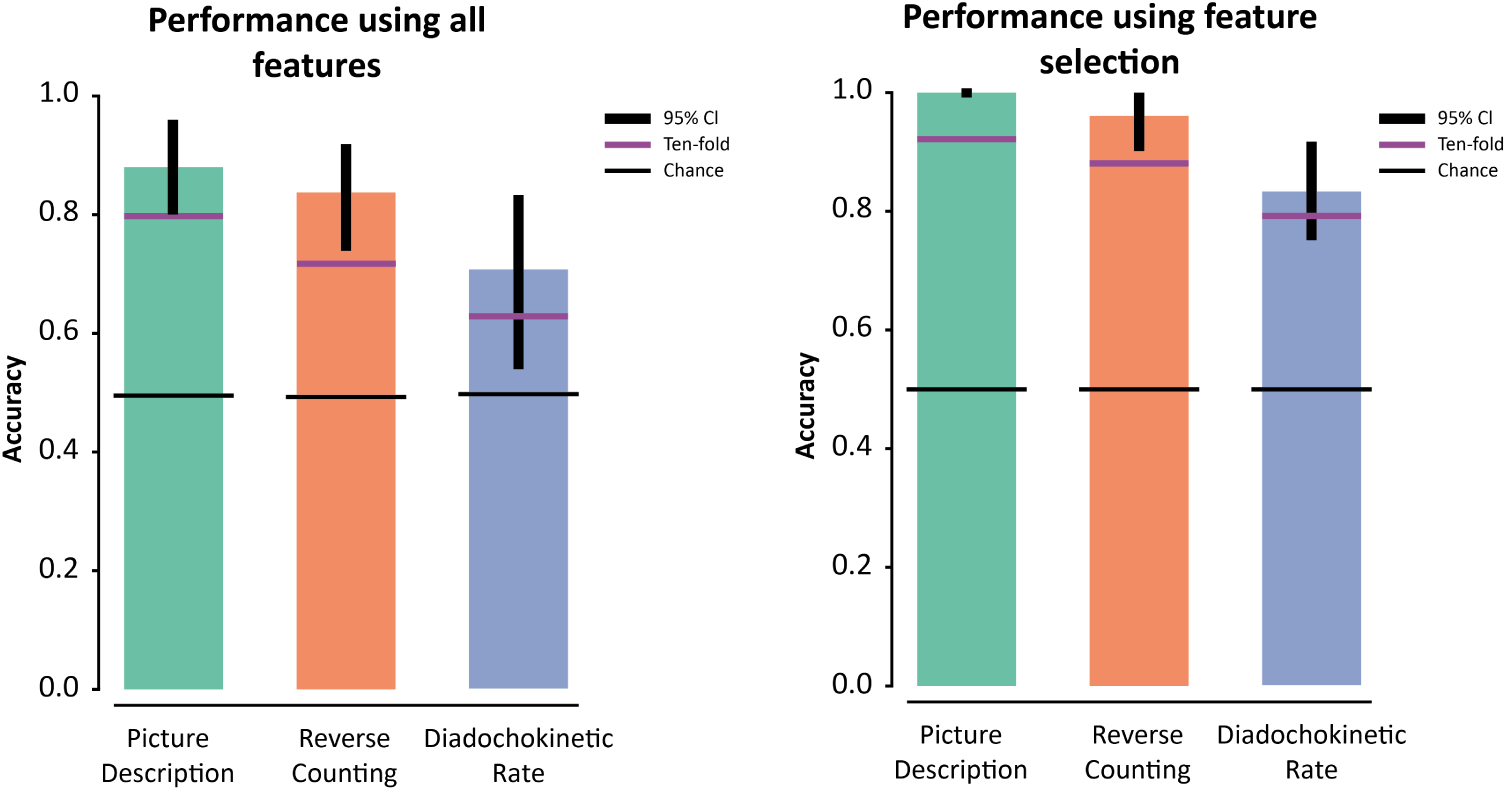
Classification performance for each task using all features (left figure) and after performing feature selection (right figure) using leave-one-subject-out cross validation. Confidence intervals at 95% are marked with black vertical lines. Chance probability is calculated for each speech task and displayed with a horizontal black line. We also indicate the accuracies obtained for ten-fold cross validation with horizontal purple lines. Results obtained with leave-one-subject-out and ten-fold cross validation surpass chance probability.

### | Impact of Audio Duration

Figure 3 shows the effects in accuracy of fixing the duration of the speech recording before feature extraction. Median accuracy value of 0.70 using 16 subjects in comparison of 0.88 using 25 subjects is achieved for recording length of 10 seconds or more. For this experiment we use exclusively the acoustic features and only one of the possible combinations of classifier and number of selected features, chosen based on the results presented on Table 2 (LR – l_1_-norm using 130 features).

**FIGURE 3.**
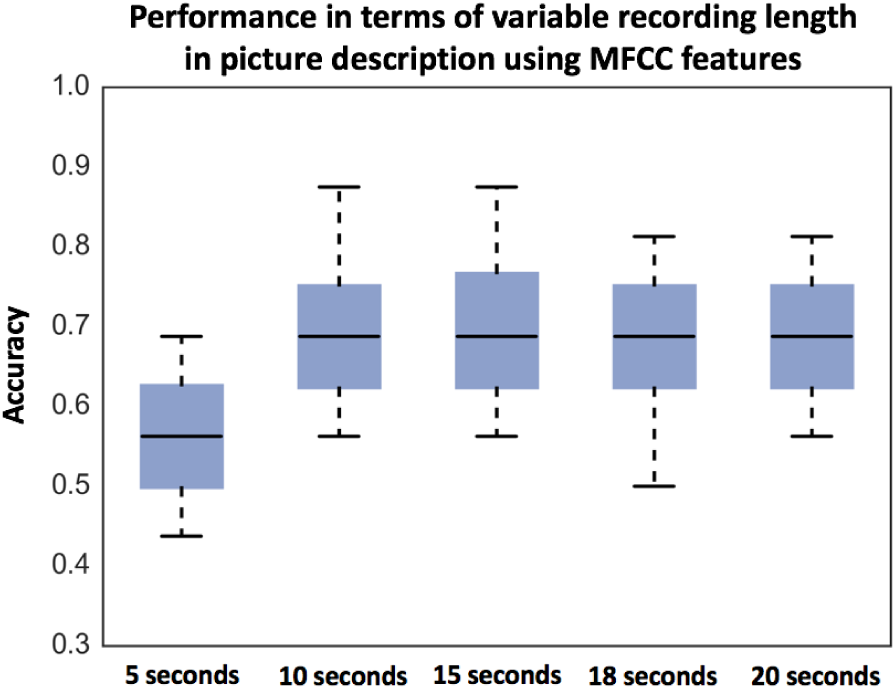
Accuracy while fixing the recording time after pause removal for 5, 10, 15, 18 and 20 seconds. Only 16 subjects were used for this analysis since we used recordings of 20 or more seconds of duration after removing pauses. The features used in this analysis were MFCC given the independence of context. Boxplots were created based on 4-fold cross validation on 100 random samples. Whiskers of boxplots show 5^th^ and 95^th^ percentiles. Median accuracy rate is stable around 0.70 for different recording lengths except for 5 seconds.

## DISCUSSION

High accuracy rates with values above chance (see Fig. 1) were achieved for all speech tasks, in particular for picture description (1.0) and reverse counting (0.96). This is consistent with the work in [45], which suggests that information extracted from running speech is better to detect PD signatures than information acquired using a diadochokinetic rate task. Furthermore, Ackermann *et al.* [33] reported that for PD subjects there may be a trade-off between amplitude of articulator movement and rate of speech affecting the results of diadochokinesis tests.

The most predominant features for the three speech tasks were MFCCs. These features can make a better characterization of the voice as they analyze it using different frequency bands. We observe in Fig. 1 that MFCCs #1 and #11 are the most significant features for reverse counting and diadochokinetic rate tasks, respectively. MFCC #1 captures information from the total frame energy, meaning that in this task the main difference between medication states is characterized by changes in the speech energy. This speech energy variation is one of the characteristics of hypokinetic dysarthria found in PDP and reported in [46]. On the other hand MFCC #11 captures information in high frequency [9.5kHz - 12.6kHz]. Recently researchers, supported by the advancements in equipment technology that captures a broader spectrum [47, 48], have shown that there is perceptually relevant information on high frequency speech, affecting speech intelligibility. Both [47, 49] concluded that high frequency speech characteristics are different in dysphonic versus control subjects, suggesting that the hoarseness characteristic of PD subjects, or in our case, the difference in hoarseness between “ON” and “OFF” states is what we are capturing with high coefficients of the MFCC. There is a report on Parkinson’s rat models [50] that finds a similar trend of what is seen on Table 1. In the rat model, the control rats had a higher maximum frequency than the dopamine-altered (reduced) rats for both the simple and frequency modulated calls.

In the picture description, the three type of features are informative and complement each other. By studying the covariation pattern shown in Fig. 1, we observe that a very high positive correlation (red lines) occurs only in “OFF” state between SF (*play*) and the other to types of features MFCC #2 and NS. This interaction observed in only one state between features help to achieve 12% improvement with respect to use only MFCC features (see Table 2). It is interesting to note on Table 1 that distance to seed words *play* and *act*, which both denote *activity* are more prominent in the ON state than the OFF state (positive sign on t-statistic).

SF are informative; however they require the context from the speaker for these to be understood. Therefore, this task needs to be designed properly. On the other hand, acoustic features are more flexible for the analysis as any part of the speech recording can be used to characterize the voice.

In addition, we also perform an extra experiment with the picture description task data to evaluate the impact of duration of the analyzed recording in the accuracy. The accuracy values are reduced from 0.88 to 0.70 as only 16 out of 25 subjects were used and 4-fold cross validation was implemented instead of leave-one-out. Nevertheless, we observe in Fig. 3 that the accuracy results are very stable when 10 or more seconds of recording are analyzed. This small duration makes feasible the implementation in mobile applications that can be used as part of a daily task to monitor PD subjects.

The experimental design balanced the visit order (“ON” in first visit 50% of the cases) to avoid interference with the classification results and the visit order. Nevertheless we tested classification using the visit order as the class. For syllable repetition where the task is exactly the same in both visits the accuracy rate is 0.52 (just acoustic features) and 0.50 (acoustic plus prosody features), thus we can assume that there is not much to learn from one visit to the next. For the reverse counting task, the accuracy rates are 0.68 (just acoustic features) and 0.64 acoustic plus prosody features; even though the task is not identical in both visits since the initial number is different, in both cases is a reverse counting using 3 as decrement unit, we interpret the results as the second time the task should be more familiar than the first time thus some learning may occur. Better accuracy rates are obtained for the picture description task, 0.84 both when using just acoustic features and when using acoustic and prosodic features; we can speculate on two reasons for this fact, there could be some learning effect, you are describing a picture both times or the medicine state is confounded with the picture used in each visit; for example the picture used for visit one, regardless of perceived L-DOPA state (ON/OFF) is easier (harder) to describe with respect to the picture used for the second visit. The conclusion from this experiment is that our models, which are different when classifying visit order or medicine ON/OFF state, are robust enough to overcome any learning effect or confounding effect with visit order and are able to distinguish L-DOPA state differences.

To find the possible causes for misclassification, we inspected the most frequently misclassified subjects. We observed, among the top reasons, that misclassified subjects present in their recordings high level of saturation and background noise produced by variation in the settings of the preamplifier. Further work will address these technical limitations, and potentially better performance may be achieved.

Finally, we want to mention that in our dataset, there is not much variation in UPDRS speech scores between medication states even though the subjects present plenty of variation in their speech. In fact, for 64% of the subjects the difference is 0, while for one case there is an improvement in speech score in OFF state. When we compared the correlation between the UPDRS total score difference and the UPDRS speech score difference, we obtained a low value of R^2^ = 0.247 with p-value of 0.01. Therefore, we want to emphasize that a new quantitative metric to monitor the patient are promising to see both disease progression and the effect on the medication(s) used. Mathematical/computational analysis of speech can increase the granularity in the assessment and also avoid the human biases that results in inconsistencies between graders.

### CONCLUSIONS AND LIMITATIONS

Our study explored different speech tasks that are easy to implement as a daily routine for monitoring PDP, obtaining high accuracy rates in detecting medication states. Our best results are obtained in the picture description task, which is a type of free speech. MFCC features are well known for capturing emotions [51, 52] and we believe that this fact may helped improving the classification accuracy since subjects can express emotions while describing the picture. Overall, our accuracy results range from 83% to 100% on naturalistic speech tasks demonstrate the potential of our analyses to be used as proxy to monitor subjects on a daily basis.

Given that this study involves a small cohort of PD subjects, a large study where different sites to enroll patients are involved will be required for further validation. In addition, the study will need to include higher variability in levels of speech score as well as higher differences of score between medication states.

To the best of our knowledge, this is the first paper to combine acoustic and semantic features of speech to monitor PD medication state. This opens the real possibility for continuous, unobtrusive, remote patient state monitoring.

## ACKNOWLEDGEMENTS

We thank Stephen Heisig for technical help with data access and Rachel Ostrand with help with protocol and audio specs.

## CONFLICT TO INTEREST

Nothing to report.

## DOCUMENTATION OF AUTHOR ROLES

Research project: Conception: JJR, GAC, RN; Data provider: BKH; Statistical Analysis: RN, CA; Review and Critique: GAC; Manuscript Preparation: RN, CA; Writing of the first draft: RN, CA; Review and Critique: GAC, JJR.

